# Isolation and primary culture of *Octopus vulgaris* cells: Assessment of their proliferative capacity

**DOI:** 10.1101/2025.04.07.647602

**Authors:** A. Galindo, R. Coello, I. Baños, E. Ramos-Trujillo, A. E. Morales, J. M. Arrieta, C. Rodríguez, E. Almansa

## Abstract

The octopus cell culture system represents a useful tool in various fields, such as biological studies and sustainable octopus production. In this paper, we describe the isolation procedure and culture protocols, metabolic activity, and the proliferative capacity assessment of the common octopus (*Octopus vulgaris*) cells, at three different life stages: embryo, paralarvae and adult. Isolation of cells was carried out with minimum essential medium (MEM) or Leibovitz L-15 medium, using 0.4 % type I collagenase for 2 (paralarvae and embryo) or 3 h (adult) at 25 °C. Isolated cells showed significant metabolic activity, demonstrating their viability for *in vitro* studies. The culture of isolated cells in MEM or L-15 supplemented with different sera (4 % fetal bovine serum (FBS), 10 % FBS and 5 % haemolymph (HEMO)) was viable for 168 h, although the survival of different cell types was variable. A detachment protocol using trypsin was also successfully developed. Finally, isolated muscle cells from both paralarval and adult specimens were marked with the mitotic indicator PHH3. Results from flow cytometry analysis showed that all developmental stages examined contained cells in mitotic phase, with embryos XII-XIII showing the highest proportion. This study provides novel guidelines for the *in vitro* culture of *O. vulgaris* cells, which are useful for the future development of cellular aquaculture, understood as the production of lab-grown meat for this species. Moreover, the results reported in this communication could contribute to reduce the use of animals for scientific purposes as well as providing a new option for human consumption of cephalopods.

## 1. Introduction

Cells cultured *in vitro* represent an essential tool in multiple studies for biological and medical sciences (Goswami et al., 2022; Kitaeva et al., 2020; Rinkevich, 1999; Slanzi et al., 2020). In spite of technical difficulties such as the risk of contamination, the limited number of cells and their short lifetime (Van Der Merwe et al., 2010), *in vitro* cell culture allows a reductionist approach, diminishing the number of experimental animals for biotechnological and biomedical research (Maselli et al., 2018; Rinkevich, 1999). In this sense, the establishment of primary cell lines is useful to advance in animal production including aquaculture, and for biotechnological, and pharmaceutical industries purposes. Invertebrates, which include more than 95 % of the animal species (Eisenhauer and Hines, 2021), could also be considered as a valuable source for cell culture practices (Rinkevich, 1999). In particular, some species such as *Drosophila melanogaster* and *Caenorhabditis elegans* have several established cell lines from tissues (Christensen et al., 2002; González et al., 2011). The culture of the whole animal and of isolated cells from *C. elegans* has also emerged as an essential tool for developing future therapies aimed at halting the loss of neuronal structure and function (Caldero-Escudero and Romero-Sanz, 2024; Zhang and Kuhn, 2013). However, one of the main challenges, especially in marine invertebrates, is the limited understanding of cell physiology, along with the slow proliferation rates and the need for optimized culture media (Van Der Merwe et al., 2010). Marine invertebrate cell cultures are of particular interest both as model organisms and for their biotechnological potential, given their ability to produce a wide range of bioactive compounds (Romano et al., 2022). Despite this relevance, only a few reports of primary cell cultures from *Porifera*, *Cnidaria*, *Crustacea*, *Mollusca*, *Echinodermata*, and *Urochordata* have been published (Rinkevich, 1999; Blues Project, 2024).

In molluscs, cell cultures have been determinant to the basic knowledge of complex physiological processes, generating information impossible to be obtained from entire animals (Yoshino et al., 2013). Thus, neuronal cell culture from gastropods helped to understand the complex neuronal networks in these animals, while primary cell cultures from bivalve and gastropod species have been useful in biomonitoring of environmental contaminants, among others (Yoshino et al., 2013). In addition, significant advances in muscle and neuronal differentiation in *Mytilus trossulus* have been attained (Dyachuk, 2013; Maiorova and Odintsova, 2016; Odintsova et al., 2010, 2000). However, a substantial portion of the relevant research remains unpublished in peer-reviewed journals, and is only available in master’s or doctoral theses, conference proceedings, and specialised books (Domart-Coulon and Blanchoud, 2022).

Within molluscs, the cephalopods are a group that has received little attention in cell culture despite their advances and contribution in several fields of knowledge such as neurobiology, development, physiology, evolutionary genomics, and biomechanics models among many others (Albertin and Simakov, 2020; Deryckere et al., 2020; Elagoz et al., 2024; Flash and Zullo, 2023; Prado-Álvarez et al., 2022). In this regard, only some advances regarding adult cell isolation (Nesher et al., 2019), neurons primary cell culture (Maselli et al., 2018) or primary cell culture from several tissues and ages (Kim et al., 2025) have been addressed. For all of the aforementioned reasons, developing new primary cell cultures of cephalopods could pave the way to improve our understanding in the knowledge areas above mentioned.

Cephalopods are one of the most important fisheries resources (Abad et al., 2023; Ainsworth et al., 2023) and some cephalopods groups such as octopods are facing an increasing demand which, together with the difficulty of managing their wild stocks, puts the sustainability of these fisheries at risk (EUMOFA, 2021; Pita et al., 2021; Rosa et al., 2024; Sauer et al., 2021). For this reason, alternative production ways such as aquaculture has been proposed to avoid additional pressure in the wild octopus population (FAO, 2024; Sauer et al., 2021), a task which is in progress for some species such as *Octopus vulgaris* and *Octopus maya* (Almansa et al., 2025; García-Fernández et al., 2019; Márquez et al., 2023; Reis et al., 2021; Tur et al., 2020, among others). So far, octopus farming has not yet been proved as ethical or sustainable (Jacquet et al., 2019), thus further research into more sustainable alternative modes of aquaculture is necessary (Almansa et al., 2025; Gleadall et al., 2025). In this regard, primary cell culture of octopus could open the door to potential biotechnological applications including research and the production of lab-grown animal meat to reduce the use of animals for human consumption. Octopus seems to be good candidates to obtain muscle cell lines given the high growth and regenerative capacity of their arms, which can be due to a relevant multipotency, totipotency, and neoplasia activities that maintain this regenerative capacity (Rinkevich, 1999). This approach could potentially lead to the development of “cellular aquaculture” for this species.

The main objective of this work was to develop protocols for cell isolation and culture at different developmental stages (embryo, paralarvae and adult) of *O. vulgaris,* that allow *in vitro* testing to be carried out. Additionally, the proliferative capacity of the isolated cells in the different stages was also studied to assess the potential development of these novel octopus cell lines.

## 2. Material and Methods

### 2.1. Ethical Statement

All the experimental procedures were carried out at the Centro Oceanográfico de Canarias (Instituto Español de Oceanografía; IEO-CSIC). Animal experiments were performed according to the Spanish legislation RD53/2013 within the framework of the European Union directive on animal welfare (Directive 2010/63/EU) and approved by the Ethics and Animal Welfare Committee of La Laguna University and competent authority of the regional government of the Canary Islands-Spain (CEIBA 1377-2023 and CEIBA 1610-2024). Furthermore, for the care and welfare of cephalopods during experimentation, we followed the guidelines by Fiorito et al. (2015) and Ponte et al. (2023).

### 2.2. Broodstock and sampling

The *O. vulgaris* adult individuals were captured by professional artisanal fishermen in the northwest coast of Tenerife, Canary Islands, Spain (28^◦^30′N, 16^◦^12′W). Upon arrival at the culture facilities of the Centro Oceanográfico de Canarias, they were weight and individually acclimated in 4000 L rectangular fibreglass tanks. The tanks were maintained under a natural photoperiod for the geographical location (10L:14D). Temperature and salinity were 22.61 ± 1.11 °C and 36.8 ± 0.1 PSU, respectively. Temperature was measured with a Tinytag Plus 2 thermometer (TGP-4020; Gemini Data Loggers Ltd., Tynitag West Sussex, United Kingdom) and salinity with a Refractometer S/Mill-E (ATAGO). About 50 % of each tank surface was covered with a green shading net. The water flow of the rearing tank was 5 L min^-1^. A mixture of frozen food, consisting of squid (*Loligo opalescens*) and crab (*Callinectes sapidus*), was fed *ad libitum* to the broodstock daily. PVC pipes and clay pots were placed inside the tanks to provide 2-3 dens for each specimen. The presence of eggs was verified on a weekly basis to avoid disturbing the breeders. When an egg mass was observed, the female with the egg mass were removed and placed in a separate 1000 L tank. Moreover, the tank filter was changed to a 500 μm mesh and the paralarvae hatched after the incubation period were moved to experimental tanks on the same day of hatching (Reis et al., 2021).

For this study, eggs from stages VI-VII and XII-XIII were selected based on Deryckere (2020), and availability of eggs. Embryos were removed from the chorion using a scalpel, and pools of 20-25 individuals were taken for each analysis. In the case of paralarvae, they were anaesthetised in 2 % (v/v) ethanol in seawater before sacrifice by mechanical destruction of the brain. After that, the arms were carefully dissected from each specimen, and arm samples from groups of 20-25 individuals were pooled. Finally, 3 adults per assay were anaesthetised using 2 % (v/v) ethanol in seawater before a 1 cm sample of the arm was excised.

### 2.3. Cell isolation from octopus embryo, paralarval and adult samples

Cell isolation was developed following Nesher et al. (2019) with small modifications. Two media were utilised depending on the experiment, a minimum essential medium (MEM; Gibco, Thermo Fisher Scientific, Waltham, Massachusetts, USA) or Leibovitz L-15 medium (Thermo Scientific). Both were modified with 400 mM NaCl, 10 mM KCl, 15 mM HEPES and 1 % (v/v) antibiotic and antimycotic solution (Merck, Darmstadt, Germany), pH 7.2. The media were then filtered under sterile conditions using a 0.22 μm pore size membrane filter.

Preliminary trials were carried out to select collagenase concentration, time, and temperature for the different samples taken (whole embryos or paralarval and adult arms samples). Enzymatic dissociation was assessed by incubating tissues in 0.4% (w/v) type I collagenase from *Clostridium histolyticum* (Merck), for 2 (embryos and paralarval muscle) or 3 h (adult muscle) at 25 °C followed by mechanical dissociation of any remaining tissues using a micropipette. The resultant cell suspensions were filtered through a 60 μm nylon mesh with the corresponding media. Cells were collected by centrifugation (Eppendorf 5425 R, Hamburg, Germany) at 200 x g for 5 min, washed and re-suspended in the medium. A sample was taken to assess the viability of cells by using the trypan blue exclusion test (> 90 % of viability in all cases).

For the metabolic activity assay, cell samples of dissociated muscle from paralarval and adult arms were frozen with liquid nitrogen and stored at −80 °C until the analysis.

### 2.4. Metabolic activity

The metabolic activity of both the intact paralarval and adult arms and isolated cells attained from paralarval and adult arm-muscle samples was enzymatically determined to assess the viability of the cells. All enzymes were measured spectrophotometrically in duplicate at 25 °C using a PowerWaveX microplate scanning spectrophotometer (Bio-Tek Instruments, Inc., Winooski, Vermont, USA).

Samples were homogenised in triplicates using four volumes of ice-cold 100 mM Tris-HCl buffer with 0.1 mM EDTA and 0.1 % (v/v) Triton X-100 (pH 7.8). After centrifugation at 30,000 × g for 30 min at 4 °C, supernatants were aliquoted and stored at −80 °C for further enzyme assays. Three intermediary metabolism enzymes were studied: lactate dehydrogenase (LDH, EC 1.1.1.27), citrate synthase (CS; EC 4.1.3.7), and glutamate oxaloacetate transaminase (GOT; EC 2.6.1.1).

The assay conditions in a final volume of 0.2 mL were as follow (Morales et al., 2017): LDH: 50 mM imidazole-HCl buffer (pH 7.4), 0.2 mM NADH, and 2.5 mM pyruvate as substrate.

CS: 50 mM imidazole-HCl buffer (pH 8), 0.1 mM DTNB, 0.2 mM acetyl CoA, and 0.2 mM oxaloacetic acid as substrate.

GOT: 50 mM imidazole-HCl buffer (pH 7.4), 10 mM α-ketoglutarate, 0.3 mM NADH, 0.05 mM pyridoxal phosphate, 3 UI mL^-1^ MDH, and 25 mM L-aspartate as substrate.

The soluble protein concentration of homogenates was quantified by the method of Bradford (1976), using bovine serum albumin as standard. Specific activity of enzymes was expressed as mU mg protein^-1^. One unit (U) of activity was defined as the amount of enzyme required to transform 1 μmol of substrate per minute under the above assay conditions. The millimolar extinction coefficients for NADH and DTNB were 6.22 (Ԑ_340_) and 13.6 (Ԑ_412_) mM^-1^ cm^-1^, respectively.

### 2.5. Cell culture in different media and supplements

Cell culture was developed following Odintsova et al. (2010) with small modifications. Thus, two experiments were conducted on isolated cells from paralarval arms. In the first one, either MEM or L-15 was tested as the culture medium. Thus, cell extracts, together with 1 mL of media were seeded in multiwells plates (6 wells per treatment) coated with poly-L-lysine, and incubated at 18 °C. Half of the volume in each well was changed every 48-72 h. Five random fields in each well were photographed at 20x using an epi-fluorescence microscopy (Zeiss, Jena, Germany), and a camera (Hamamatsu ImagEM, Shizuoka, Japan), 24 and 144 h after the seeding. Images were analysed using the AxioVision 4.8. software.

In the second experiment, and in order to define the best nutritional supplement for paralarval cell culture, 4 % fetal bovine serum (FBS; Thermo Scientific), 10 % FBS and 5 % octopus’ haemolymph were tested.

Haemolymph was directly collected from the branchial blood vessel using a sterilised syringe (Malham et al., 1998) from two adult octopuses, which were previously anaesthetised. Haemolymph was diluted (1:1) in a marine anticoagulant solution containing 0.1 M glucose, 15 mM sodium citrate, 13 mM citric acid, 10 mM EDTA and 0.45 M NaCl (pH 7.0 and 1000 mOsm) (Barcia et al., 1999). The resulting solution was centrifuged for 10 min at 800 x g and 4 °C. Supernatant was sterilised by filtering through a 0.20 μm pore size filter, and stored at −20 °C until its utilisation.

Isolated cells from paralarval arms were seeded in multiwell plates (4 wells per treatment) coated with poly-L-lysine, adding 1 mL of MEM, together with the corresponding supplement, and cultivated at 18 °C. Half of the medium of each well was changed every 48-72 h. Five random fields from each well were photographed at 20x, after 72 and 168 h from seeding, as previously explained. In both experiments, survival of cells was calculated by dividing the number of cells at the end of the experiment (144 or 168 h) between the number of cells at the beginning of the experiment (24 or 72 h, respectively).

### 2.6. Cell attachment with different coating agents and cell detach

In order to improve cell attachment, coating agents such as Geltrex (Thermo Scientific) and type I collagen (Merck) were assayed. Cells from embryos and paralarval arms were isolated as previously explained. Cell extracts and 1 mL of medium were seeded in multiwells plates (3 wells per treatment) coated with either Geltrex or type I collagen, and incubated at 18 °C for 48 h. Subsequently, a procedure using different conditions to detach the cells was developed. A first trial was conducted using three different solutions: calcium and magnesium-free Hanks’ Balanced Salt Solution (HBSS) containing 1.75 % (w/v) NaCl, 9.69 mM HEPES, 1.73 mM NaHCO₃, and 5 mM EDTA; a 0.4 % (w/v) collagenase solution; and a 4 % (w/v) trypsin solution. These treatments were applied to paralarval arms and embryo cells seeded on Geltrex and type I collagen. Since trypsin showed the best results, a second trial was carried out to determine the optimal detachment conditions. For this purpose, the detachment of embryo cells cultured on type I collagen was assessed using 4% trypsin under different conditions: two incubation temperatures (18 and 37 °C), different trypsin exposure times (1 and 3 minutes), trypsin temperatures (room temperature and 37 °C, respectively), and the presence or absence of a cell scraper (Table 1).

**Table 1.** Different combinations used to test trypsin as detaching agent of embryo cells attached to type I collagen.

| Culture cell temperature | Trypsin time | Trypsin temperature | Cell scraper |
| --- | --- | --- | --- |
| 18 °C | 1 min | Room temperature | No |
| 18 °C | 3 min | 37 °C | No |
| 18 °C | 1 min | Room temperature | Yes |
| 18 °C | 3 min | 37 °C | Yes |
| 37 °C | 3 min | 37 °C | Yes |

### 2.7. Cellular characterization of embryo and paralarvae using immunocytochemistry

In order to characterise the type of cells obtained from the whole embryo, and from paralarval pooled arms, and to test the existence of mitotic cells, immunocytochemical assays were developed using the monoclonal antibody anti-phospho-Histone H3 (PHH3) as a mitotic marker. Cells were seeded in multiwell dishes on glass coverslips coated with poly-L-lysine and cultivated at 18 °C in MEM. After 72 h, cells were washed with phosphate buffered saline (PBS), fixed with 4 % (w/v) paraformaldehyde (PFA) in PBS for 15 min, and washed again with Tween 20 in PBS (PBT) for 15 min under shaking. Wells were then incubated with 5 % (v/v) of goat serum in PBT for 1 h and shaking, and later with the primary antibody PHH3 (pSer28) (Merck) 1:600 (v/v) in PBT and 2.5 % (v/v) goat serum for 2 h at room temperature. Samples were washed again three times with PBT for 10 min in the dark and under shaking, and subsequently incubated with the secondary goat antibody (anti-rat) (Alexa Fluor® 488; Thermo Fisher Scientific) 1:500 (v/v) in PBT for 1 h at the dark and shaking. Samples were then washed 3 times with PBT for 10 minutes with shaking. Finally, samples were mounted with Fluoroshield containing DAPI (Thermo Scientific) and imaged with an epi-fluorescence microscopy (Zeiss), using a camera (Hamamatsu ImagEM) and AxioVision 4.8. software.

### 2.8. Cell cycle profile by flow cytometry

Cell cycle profile was determined in isolated cells from embryos and paralarvae and from adult arms (3 replicates per developmental stage). Simultaneous staining of DNA content with propidium iodide (PI; Merck) and the mitotic marker PHH3 with Alexa Fluor®488 (Thermo Fisher Scientific) was developed, using a hypotonic buffer to permeabilize cells (Shen et al., 2017). Briefly, cells were resuspended in Dulbecco’s phosphate buffered saline (DPBS). Then, they were centrifuged at 200 x g for 5 min and resuspended in a hypotonic buffer containing deionized water, 0.1 % (w/v) sodium citrate, 0.1 % (v/v) Triton X-100 and 50 μg/mL PI. After adding 10 µL of Alexa Fluor® 488, samples were incubated for 20 min in the dark at room temperature and analyzed on a CytoFLEX S flow cytometer (Beckman Coulter, Inc.) using CytExpert software (version 2.4). The red fluorescence of PI was measured through a 585/42 nm band-pass (BP) filter, while the green fluorescence of Alexa Fluor®488 was detected with a 525/40 nm BP filter. G_1_, S and G_2_/M phases were differentiated on the basis of DNA content histograms; whereas M phase cells were clearly identified by DNA content versus Alexa Fluor®488 dot plots (Figure 1A-B). Residual debris was excluded from the analysis.

**Figure 1.**
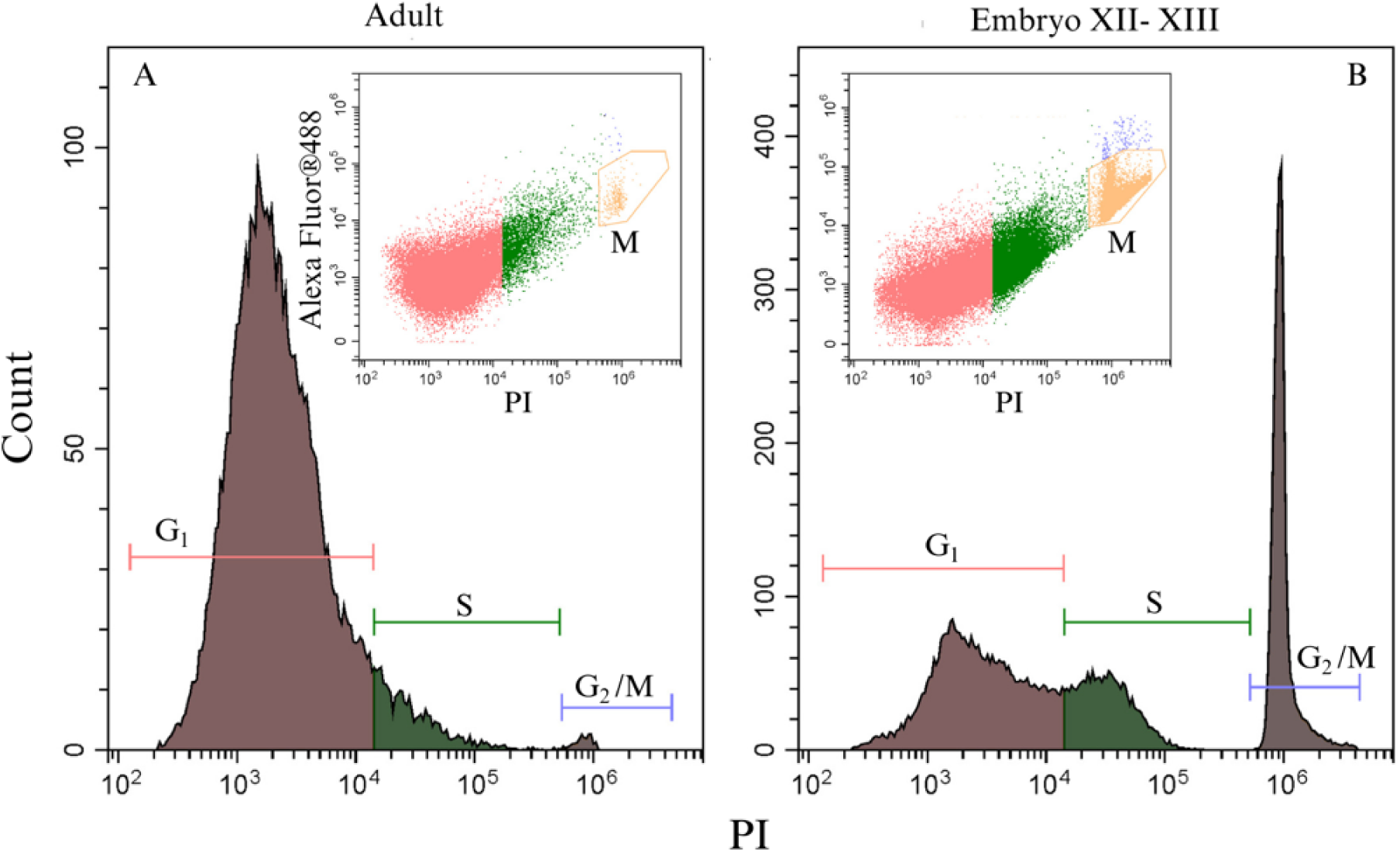
Histogram of DNA content (PI) for G_1_, S and G_2_/M phases in adult muscle cells (A) and embryo XII-XIII cells (B). (*Inner panel*) Identification of M phase by dot plots of DNA content (PI) versus Alexa Fluor®488 stained cells.

### 2.9. Statistical analysis

Assumptions regarding normality and homoscedasticity of data were assessed, and arcsine variance-stabilising transformation was applied when necessary. Significant differences were determined using one-way ANOVA followed by a Tukey HSD post-hoc test. For data lacking homoscedasticity, a Welch test followed by the Dunnett T3 test was employed. In cases of non-normal distribution, the Kruskall-Wallis nonparametric test was used, followed by pairwise comparison Mann–Whitney test and a Bonferroni correction. Additionally, Student’s t-test or Mann-Whitney tests were utilised depending on whether the data followed a normal or non-normal distribution.

Data are presented as means ± standard deviation (SD), with statistical significance set at p<0.05. Statistical analysis was carried out with IBM SPSS Statistics 25.0 software package (IBM Corp., New York, USA) for Windows.

## 3. Results

### 3.1. Metabolic activity of isolated cells

To assess the metabolic activity and viability of cells isolated from paralarval and adults arms, several intermediary metabolism enzymes such as CS, GOT and LDH, were measured and compared with values obtained in the corresponding intact tissues (Table 2). Overall, the results showed lower but significant activity in isolated cells compared with their original tissues. Thus, CS, GOT and LDH activities were higher in adult muscle than in adult isolated muscle cells. Similarly, GOT activity was higher in paralarval intact tissue, whereas CS and LDH activities showed similar values in intact tissue and isolated cells (Table 2).

**Table 2.** Enzymatic activity (mU mg protein^-1^) of citrate synthase (CS), glutamate oxaloacetate transaminase (GOT) and lactate dehydrogenase (LDH) in both adult and paralarval intact tissues, as well as their corresponding isolated cells.

| Enzyme | Adult tissue | Adult cells | Paralarval tissue | Paralarval cells |
| --- | --- | --- | --- | --- |
| CS | 36.60 ± 4.29 | 24.47 ± 5.83 * | 32.73 ± 5.08 | 23.83 ± 5.15 |
| GOT | 346.43 ± 18.18 | 154.06 ± 23.59 * | 254.65 ± 22.98 | 93.85 ± 14.92 * |
| LDH | 95.46 ± 15.66 | 57.83 ± 14.92 * | 81.03 ± 12.38 | 54.79 ± 12.80 |

Results are presented as means ± SD (3 replicates per sample type). \* Represent significant differences (Student's t-test) between intact tissue and isolated cells for the same enzyme (p<0.05).

### 3.2. Optimization of octopus paralarval cell culture

To optimise octopus paralarval cell culture, we compared different culture media and assessed the relative abundance of the cell types found and their survival. Main cell populations (> 80 %) were defined as type I and II. Type I (in red) included round-shaped cells, and type II (in green) were defined as round-shaped cells with a dark protrusion (Figure 2).

**Figure 2.**
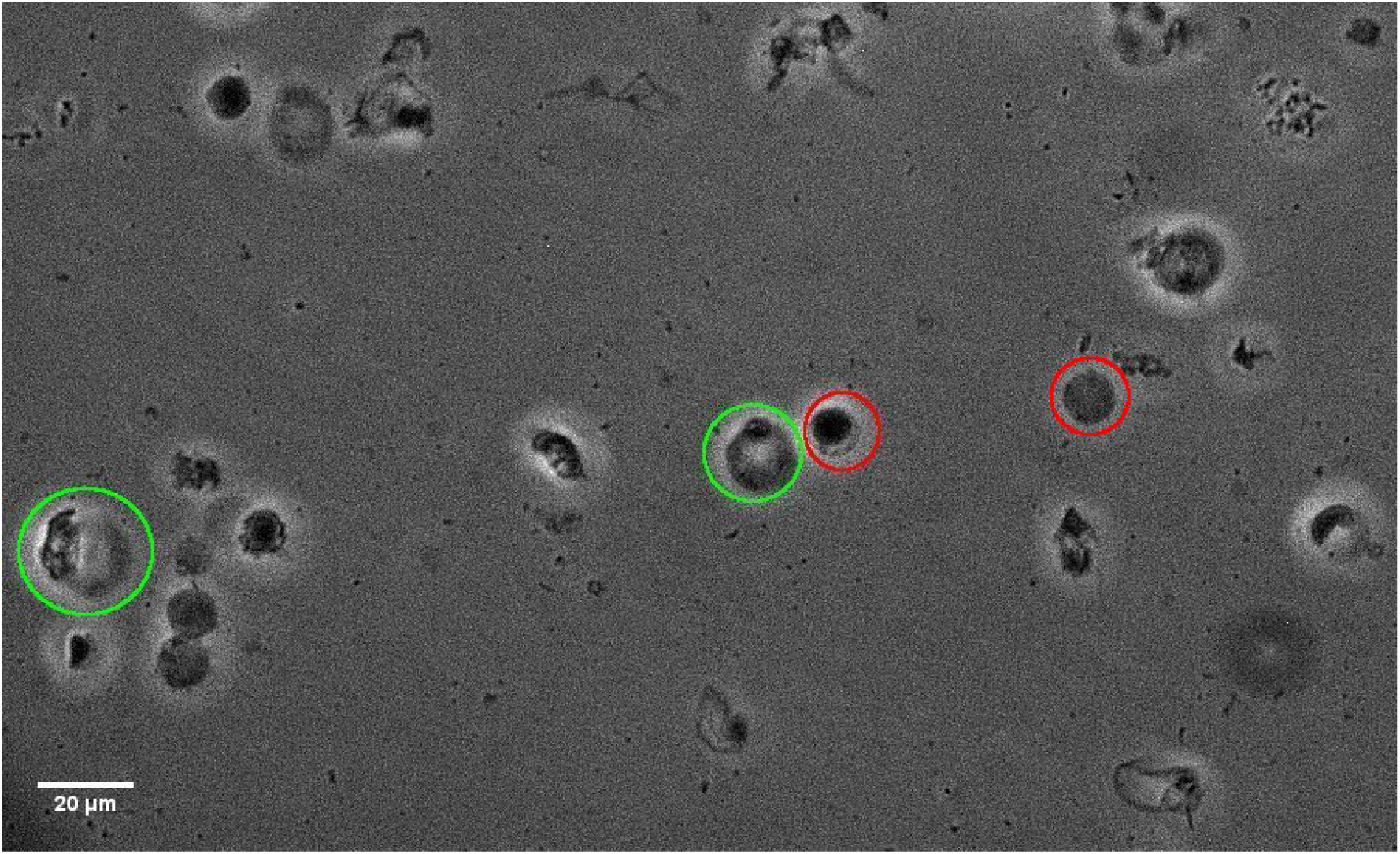
Cell types detected in paralarval cell culture at 20x. Circle red includes type I cells; circle green includes type II cells.

After 24 h, type I cells encompassed ∼60 % regardless of the medium used, with similar values (∼57-59 %) after 144 h of culture. On the other hand, L-15 showed a higher proportion of type II cells at 24 h. Overall, proportions of type II cells increased after 144 h of culture (Table 3).

**Table 3.**
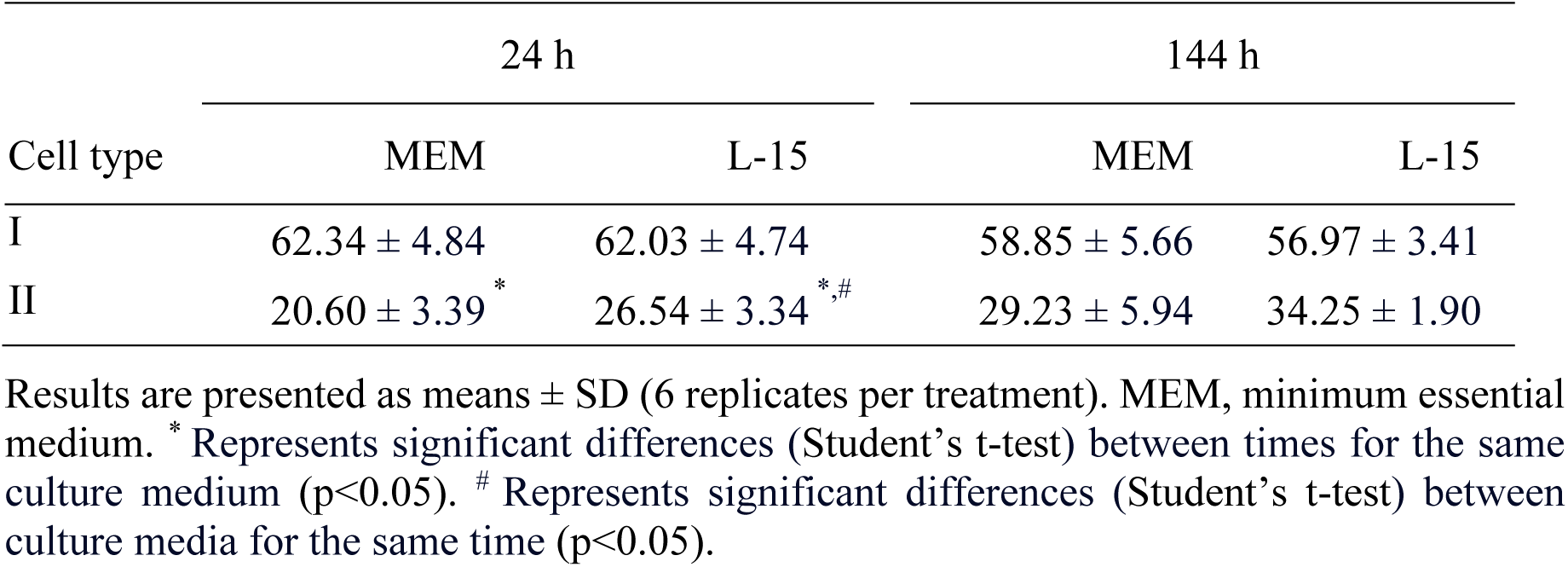
Relative abundance (%) of type I and type II cells detected in paralarval cell culture after 24 and 144 h using different culture media.

| Cell type | 24 h |  | 144 h |  |
| --- | --- | --- | --- | --- |
|  | MEM | L-15 | MEM | L-15 |
| I | 62.34 ± 4.84 | 62.03 ± 4.74 | 58.85 ± 5.66 | 56.97 ± 3.41 |
| II | 20.60 ± 3.39 * | 26.54 ± 3.34 *,# | 29.23 ± 5.94 | 34.25 ± 1.90 |

Results are presented as means ± SD (6 replicates per treatment). MEM, minimum essential medium. \* Represents significant differences (Student's t-test) between times for the same culture medium (p<0.05). # Represents significant differences (Student's t-test) between culture media for the same time (p<0.05).

Recorded survival values (between 144 and 24 h) were higher in MEM, with ∼62 %, ∼92% and ∼65 % of type I, II and total cells, respectively (Figure 3), although observed differences were not significant.

**Figure 3.**
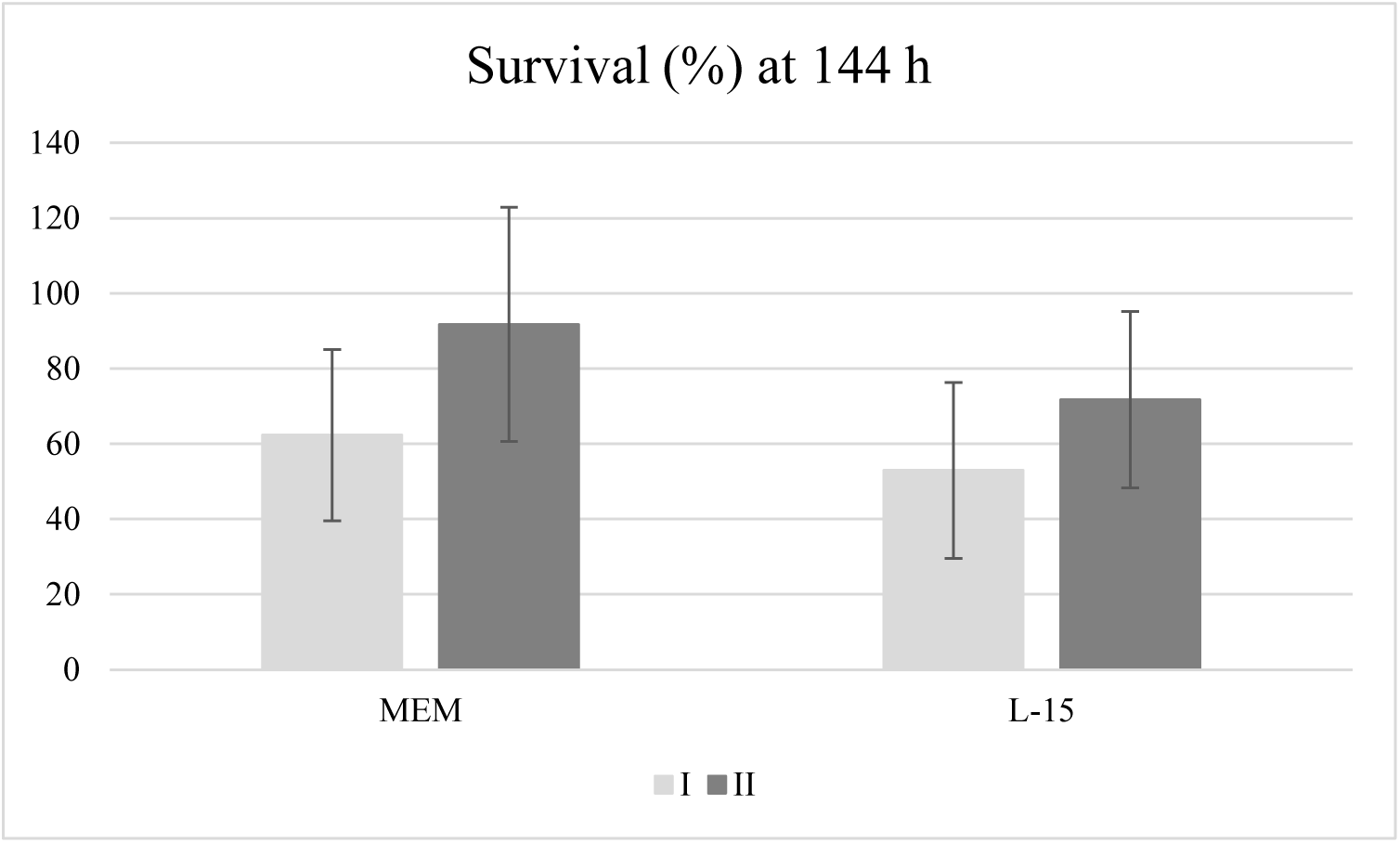
Survival (%) of paralarval cells after 144 h of culture in the different culture media. Results are presented as means ± SD (6 replicates per treatment). MEM, minimum essential medium

Similarly to the previous tests, paralarval cells were used to test different supplements such as FBS at 4 and 10 % as well as HEMO at 5 %. Results showed that these supplementary treatments did not have any effect after 72 h in any cell type. However, a positive effect was observed after longer incubation times: after 168 h, 4 % FBS favoured type I cell (∼78 %), followed by 10 % FBS (∼68 %) and 5 % HEMO (∼59 %), while 10% FBS and 5 % HEMO were the best treatments to maintain type II cells (28-34 %) compared to 4 % FBS (∼18 %) (Table 4).

**Table 4.** Relative abundance (%) of type I and type II cells detected in paralarval cell culture after 72 and 168 h using different nutritional supplements.

| Cell type | 72 h |  |  | 168 h |  |  |
| --- | --- | --- | --- | --- | --- | --- |
|  | 4 % FBS | 10 % FBS | 5 % HEMO | 4 % FBS | 10 % FBS | 5 % HEMO |
| I | 66.68 ± 2.05 | 61.82 ± 3.15 | 65.72 ± 4.73 | 77.83 ± 7.71<sup>b</sup> | 67.69 ± 7.61<sup>ab</sup> | 59.42 ± 3.35<sup>a</sup> |
| II | 30.13 ± 2.43 | 32.07 ± 3.65 | 28.18 ± 3.62 | 17.98 ± 5.45<sup>a</sup> | 27.85 ± 5.48<sup>b</sup> | 34.17 ± 3.51<sup>b</sup> |

Results are presented as means ± SD (4 replicates per treatment). FBS, fetal bovine serum; HEMO, octopus haemolymph. <sup>a,b</sup> Represent significant differences (one-way ANOVA) between supplements for the same time (p<0.05).

Finally, survival (between 168 and 72 h) of type I and total cells were similar regardless of the supplement used. On the other hand, 10 % FBS increased the survival of type II cells at 168 h, followed by 5 % HEMO and 4 % FBS (Figure 4).

**Figure 4.**
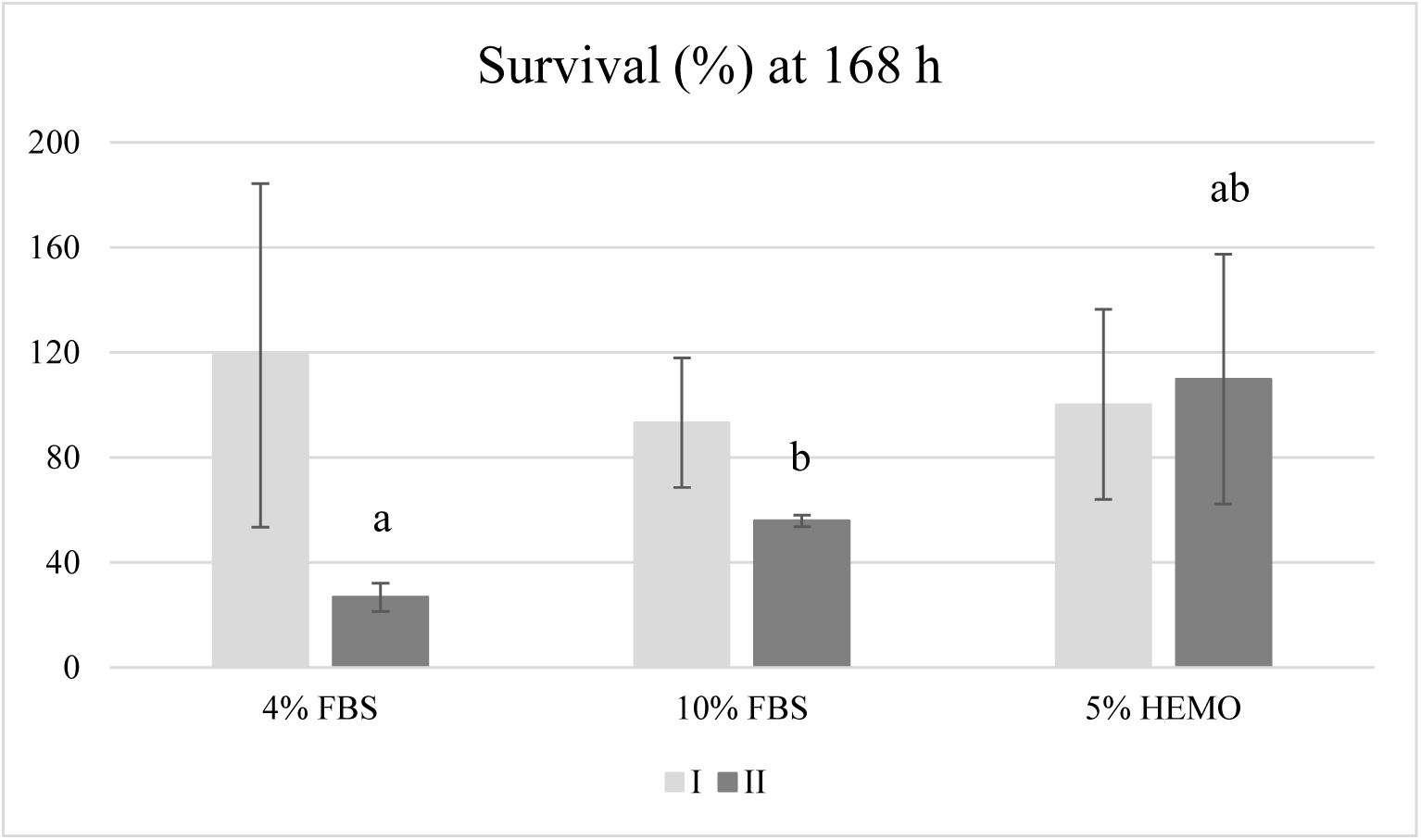
Survival (%) of paralarval cells after 168h of culture with the different nutritional supplements. Results are presented as means ± SD (4 replicates per treatment). Different letters represent significant differences (one-way ANOVA) between supplements for the same cell type (p<0.05).

### 3.3. Cell attachment and detachment

Cell extracts from embryo, or paralarval and adult arm-muscle were highly attached to both Geltrex and collagen coating agents after 48 h of seeding, with both coating agents showing similar results.

Regarding the detachment of cells from the support, neither HBSS-EDTA nor collagenase showed a proper detachment of the cells, independently of their developmental stage or the coating agent used. On the other hand, trypsin treatment showed the most promising results (> 90% of cell recovery) using a cell culture temperature of 18 °C, and letting the trypsin react for 3 min at 37 °C.

### 3.4. Proliferative capacity of the isolated cells

To achieve this objective, the first step was an immunocytochemistry characterization of potential stem cells by PHH3, a specific marker for mitosis. Both paralarval and adult arm-muscle cells exhibited the presence of mitotic cells (Figure 5A-B).

**Figure 5.**
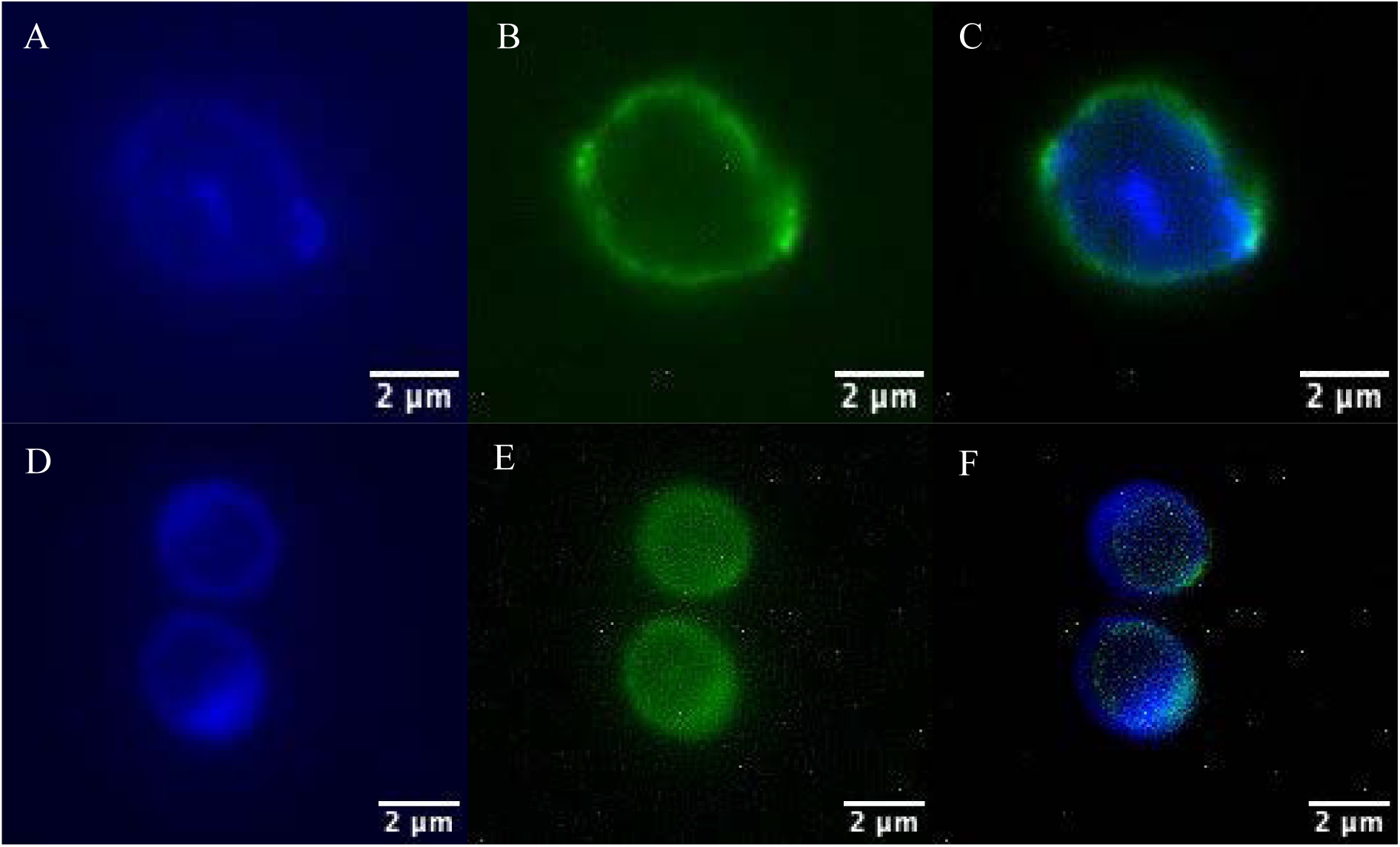
Immunocytochemistry images at 100x. Paralarval cell with DAPI (blue, A), PHH3 (green, B) and DAPI and PHH3 (C). Adult cells with DAPI (D), PHH3 (E) and DAPI and PHH3 (F).

A flow cytometry test was subsequently carried out to characterise the cell cycle phase of cells obtained at each developmental stage (Table 5). Results showed that embryo XII-XIII had the highest proportions of cells in M phase (∼23 %), followed by paralarvae (∼16%), embryo VI-VII (∼9 %) and adult muscle (∼1.4 %). Embryo XII-XIII also showed the highest percentages of cells in S phase, with 20 %. On the other hand, adult showed the highest proportions of cells in G1 phase (∼92 %), while G2 was more prominent in paralarvae (∼74 %).

**Table 5.** Relative abundance (%) of cells in each cell cycle phase at the different developmental stages.

| Phase | Embryo VI-VII | Embryo XII-XIII | Paralarvae | Adult |
| --- | --- | --- | --- | --- |
| G<sub>1</sub> | 83.20 ± 7.57<sup>bc</sup> | 56.96 ± 8.01<sup>a</sup> | 74.12 ± 3.84<sup>b</sup> | 91.98 ± 1.28<sup>c</sup> |
| G<sub>2</sub> | 0.12 ± 0.09<sup>a</sup> | 0.25 ± 0.04<sup>ab</sup> | 0.41 ± 0.08<sup>b</sup> | nd |
| S | 7.59 ± 1.71<sup>a</sup> | 20.21 ± 3.65<sup>b</sup> | 9.20 ± 0.88<sup>a</sup> | 6.60 ± 1.30<sup>a</sup> |
| M | 9.09 ± 5.83<sup>b</sup> | 22.58 ± 7.67<sup>c</sup> | 16.27 ± 3.01<sup>bc</sup> | 1.43 ± 0.65<sup>a</sup> |

Results are presented as means ± SD (3 replicates per developmental stage). Different letters in superscript within the same row represent significant differences (one-way ANOVA) between developmental stages (p<0.05).

## 4. Discussion

### 4.1. Cell isolation

Primary cell cultures are initiated using explants (tissue fragmentation) or by tissue dissociation to obtain isolated cells (Hendijani, 2017). Cell isolation can be achieved by disrupting extracellular matrix proteins with proteolytic enzymes such as trypsin or collagenase (Mótyán et al., 2013; Van Der Merwe et al., 2010). To characterise the potential for cell culture and proliferation throughout the entire developmental process of *O. vulgaris*, cell isolation from embryos, paralarvae, and adults was successfully assessed. The final protocol was defined after preliminary trials using experimental conditions similar to other studies. Thus, while Nesher et al. (2019) used a dissociation temperature of 25-30 °C in *O. vulgaris* muscle, our preliminary assays showed that temperatures of 25 °C were less effective than 37 °C, although the latter caused cellular damage. Therefore, a temperature of 25 °C was set in our experiments. Different dissociation enzymes generally used in mammalian cell culture were also tested. While papain showed better results to dissociate neural tissue from *O. vulgaris* than collagenase + trypsin (Maselli et al., 2018), in our trial testing collagenase, or trypsin + collagenase, showed no differences in either cell dissociation or integrity. Furthermore, Nesher et al. (2019) used 0.2 % collagenase for 6 h to dissociate muscle cells from *O. vulgaris.* However, in our study, following a protocol described by Maselli et al. (2018), the concentration of collagenase was increased to 0.4 % in order to reduce the isolation time to 2-3 h. Since cells are vulnerable to mechanical and toxic damage during dissociation, it has been previously recommended to reduce the time of dissociation (Van Der Merwe et al., 2010).

Finally, regarding the medium used along the dissociation process, both L-15 and MEM were tested to isolate the cells, showing similar cellular survival (trypan blue exclusion test). However, L-15 is more commonly used as the main medium in similar studies with both *O. vulgaris* and *M. trossulus* (Maiorova and Odintsova, 2016; Maselli et al., 2018; Nesher et al., 2019; Odintsova et al., 2010, 2000).

### 4.2. Metabolic activity

Monitoring of metabolic enzymes such as LDH is commonly used to assess cell viability (Haslam et al., 2000). LDH is involved in the energy production by the conversion of pyruvate to lactate during anaerobic glycolysis (Haslam et al., 2000). In particular, LDH, together with other enzymes, has been related to provide the energy needed to support burst swimming episodes during first days of paralarvae (Morales et al., 2017). In our study, values of LDH in all samples analysed were higher to values previously reported (Morales et al., 2017) corresponding to 0-12 day old paralarvae.

Other enzymes used as reporters, such as CS and GOT, are involved in aerobic metabolism (Seibel and Childress, 2000), with GOT providing the oxaloacetate necessary for the CS reaction (Morales et al., 2017). The activity of these two enzymes showed a direct relation. This would explain higher values of both enzymes in adult tissue, and the same trend (although not significant for CS) in paralarvae. CS values were generally lower in both tissue and cells, and GOT were slightly lower in cells and similar in tissues than those previously reported for paralarvae (Morales et al., 2017). However, the detection of those activities in isolated cells from *O. vulgaris* paralarvae and adult, together with the > 90 % of viability by trypan blue exclusion test, demonstrated the metabolic activity and viability of the isolated cells.

### 4.3. Culture media and nutritional supplements

Since different cell types need different culture conditions, the success in achieving primary cell cultures heavily depends on the culture medium used. This choice is usually a first challenge when working with a new species (Van Der Merwe et al., 2010). L-15 medium is widely used to cultivate cells, especially from marine invertebrates (Daugavet and Blinova, 2015; Maiorova and Odintsova, 2016; Maselli et al., 2018; Odintsova et al., 2010, 2000), while MEM is more commonly used in mammalian cell culture (Mukherjee et al., 2023). In our experiments, both L-15 and MEM were tested using isolated cells from paralarval arms. Different morphologies of cells could be observed, which may correspond to muscle, epithelial, or fibroblasts, among other cells from muscle and skin (Fernández-Gago et al., 2012). However, the cellular characterization was not the main objective of this study, being necessary additional studies in order to correctly identify the type of cells in each culture media. Our results showed similar survival rates, demonstrating that apparently both media are adequate to culture paralarval cells. Nevertheless, the impact of both media may differ depending on the cell type, and further research is required to elucidate this phenomenon.

Regarding the use of nutritional supplements, Maselli et al. (2018) suggested that the effect of nutritional supplements in octopus neuron culture was coating agent-dependent, with poly-L-lysine being the most effective coating agent, and poly-D-lysine requiring the use of nutritional supplements. In our experiments, poly-L-lysine was used as the coating reagent, while HEMO and FBS were used as nutritional supplements. The proportions and survival of different paralarval cell types varied depending on the nutritional supplement, indicating a supplement-dependent response.

### 4.4. Coating and detachment agents

Although poly-L-lysine has been described to favour *O. vulgaris* neuron attachment compared to poly-D-lysine (Maselli et al., 2018), low cell attachment was detected in this study during the immunocytochemical protocol. To enhance cell attachment, both type I collagen and Geltrex were tested, and both demonstrated similar improvements in the results. The use of different coating agents such as poly-L-lysine, type I collagen, carbon fibronectin or even uncoated glass coverslips has been previously described for cell cultures from larval *M. trolossus* (Maiorova and Odintsova, 2016; Odintsova et al., 2010). The coating agent may also affect cell mitotic capacity and differentiation. In *M. trolossus*, collagen has been shown to maintain mitotic capability for up to 6 months, while inhibiting myoblasts differentiation, in contrast to fibronectin (Maiorova and Odintsova, 2016). Thus, the importance of finding the appropriate cell culture conditions for each species, type of cells and assay is evident.

Detachment of cells from *M. trolossus* larvae cell culture has been previously accomplished by using a filter-sterilized Ca^2+^- and Mg^2+^-free salt solution with EDTA (Dyachuk, 2013; Odintsova et al., 2000), contrary to this study, were the detachment protocol using HBSS-EDTA and collagenase did not show a proper detachment in paralarval cells. Thus, the final protocol was set as using trypsin for 3 min at 37 °C. Trypsin is one of the most common enzymes to detach cells in cell culture techniques (Lordon et al., 2024).

### 4.5. Proliferative capacity

Cell cycle analysis helps to determine growth potential and changes in cell number in cultures (Leelavatcharamas et al., 2020). The protocol used in this work for dissociating and isolating cells from *O. vulgaris* allowed us to study the cell cycle and proliferative capacity of the different developmental stages by both immunocytochemistry and flow-cytometry, based on incorporated PHH3 *versus* DNA content in cells. Similar techniques have previously been applied to study neurogenesis in *O. vulgaris* (Di Cosmo et al., 2018). Immunocytochemistry was developed to test the viability of using mitotic markers. Subsequently, flow cytometry was conducted, providing a more accurate quantitative analysis compared to microscopy (Di Cosmo et al., 2018). Flow cytometry has been commonly used to detect the presence of cells proliferating *in vitro* (Lyons and Parish, 1994).

In the present study, the existence of proliferative cells is demonstrated in embryos as well as paralarval and adult arms, showing that embryos from stage XII-XIII had the highest proportion of mitotic cells. In fact, the presence of proliferative cells in embryos, with the stage XIII showing the highest number of proliferative cells, was also described by Deryckere (2020), who used PHH3 as a mitotic marker on the whole embryo. The use of tissues with high proliferative activity, which may be potentially due to the presence of stem-like cells, have been described as an essential condition to establish primary cell cultures. Aquatic invertebrates, particularly molluscs, display regeneration processes, indicating high cellular plasticity, cellular proliferation and possibly involvement of stem-like cells. However, existing protocols for the isolation and identification of stem-like cells are currently available for only a limited number of species (Domart-Coulon and Blanchoud, 2022). Further studies need to be carried out to confirm the existence of stem-like cells in those developmental stages.

## 5. Conclusion

In this study a protocol to isolate viable and metabolically active cells was developed from different *O. vulgaris* developmental stages. Cells were isolated by using 0.4 % collagenase for 2-3 h at 25 °C, in either MEM or L15. The effect of media and supplements used varied depending on the types of cells. Attachment of cells after 48 h was achieved using Geltrex and type I collagen as coating agents, while their detachment was accomplished with 4 % trypsin for 3 min at 37 °C. Immunocytochemical and flow cytometry assays were performed in order to assess cell proliferation. All developmental stages showed cells in the mitotic phase, but XII-XIII embryos were the most promising developmental stage for promoting cellular aquaculture. The results of this study can foster *in vitro* basic studies with *O. vulgaris* and also contribute to future production of lab-grown animal meat from this species. Both achievements would have positive bioethical consequences since they will significantly reduce the animal sacrifice.

## Acknowledgements

Dr. A. Galindo was sponsored by the Catalina Ruiz Programme, funded by Consejería de Economía, Conocimiento y Empleo, and FSE. This study has funded by the ThinkInAzul programme supported by Spanish Ministerio de Ciencia, Innovación y Universidades (PRTR-C17.I1), Next Generation (EU) program and Gobierno de Canarias (SD2218/6897). In addition, it has also been funded by the European Maritime, Fisheries and Aquaculture Fund (EMFAF) by the project AMTI-CANARIAS. Dr. C. Rodríguez is member of Instituto de Tecnologías Biomédicas de Canarias (ITB). We thank Dr. Julio Polaina (theenzymefishingcompany.tech), M. Lassnig and Dr. E. Seuntjens (Ku Leuven, Belgium) for useful suggestions and critical reading of the manuscript. We also thank Dr. José David Machado and Dr. Marcial Camacho (ULL, Spain) for helping with cell culture and immunocytochemistry assays.

## References

Abad, E., Ainsworth, G., Akselrud, C., Allcock, L., 2023. Working group on cephalopod fisheries and life history (WGCEPH; outputs from 2022 meeting). ICES Sci. Reports 5, 1–163. 10.17895/ices.pub.21976718

Ainsworth, G.B., Pita, P., Garcia Rodrigues, J., Pita, C., Roumbedakis, K., Fonseca, T., Castelo, D., Longo, C., Power, A.M., Pierce, G.J., Villasante, S., 2023. Disentangling global market drivers for cephalopods to foster transformations towards sustainable seafood systems. People Nat. 5, 508–528. 10.1002/pan3.10442

Albertin, C.B., Simakov, O., 2020. Cephalopod biology: At the intersection between genomic and organismal novelties. Annu. Rev. Anim. Biosci. 8, 71–90. 10.1146/annurev-animal-021419-083609

Almansa, E., Márquez, L., Rosas, C., Martín, M., Navarro, J., Uriarte, I., Gestal, C., Fernández-Álvarez, F., Gallardo, P., Varó, I., Farías, A., Cardenete, G., Caamal-Monsreal, C., Rodríguez-Barreto, D., Morales, A., 2025. Octopods aquaculture: Reproduction, rearing technology, nutritional physiology, welfare and health status, in: Pereira, P.L. (Ed.), Aquaculture and Living Resource Management – Volume 2. CRC Press (Taylor & Francis Group), Boca Ratón. In press.

Barcia, R., Cao, A., Arbeteta, J., Ramos-Martinez, J.I., 1999. The IL-2 receptor in hemocytes of the sea mussel *Mytilus galloprovincialis* Lmk. IUBMB Life 48, 419–423. 10.1080/713803540

Bradford, M.M., 1976. A rapid and sensitive method for quantitation of microgram quantities of protein utilizaing the principle of protein-dye binding. Anal. Biochem. 72, 248–254. 10.1016/j.cj.2017.04.003

Caldero-Escudero, E., Romero-Sanz, S., 2024. Caenorhabditis elegans como modelo animal de investigación científica. Caenorhabditis elegans as a research animal model. Clínica 29, 67–69. 10.1016/j.taap.2018.03.016

Christensen, M., Estevez, A., Yin, X., Fox, R., Morrison, R., Mcdonnell, M., Gleason, C., Miller, D.M., Strange, K., 2002. A primary culture system for functional analysis of *C. elegans* neurons and muscle cells. Neuron 33, 503–514.

Daugavet, M.A., Blinova, M.I., 2015. Culture of mussel (*Mytiuls edulis* L.) mantle cells. Cell tissue biol. 9, 233–243. 10.1134/S1990519X15030037

Deryckere, A., 2020. Neurogenesis, neural migration and differentiation in the developing octopus brain. KU LEUVEN, Science, Leuven.

Deryckere, A., Styfhals, R., Vidal, E.A.G., Almansa, E., Seuntjens, E., 2020. A practical staging atlas to study embryonic development of *Octopus vulgaris* under controlled laboratory conditions. BMC Dev. Biol. 20, 7. 10.1186/s12861-020-00212-6

Di Cosmo, A., Bertapelle, C., Porcellini, A., Polese, G., 2018. Magnitude assessment of adult neurogenesis in the *Octopus vulgaris* brain using a flow cytometry-based technique. Front. Physiol. 9, 1–10. 10.3389/fphys.2018.01050

Domart-Coulon, I., Blanchoud, S., 2022. From primary cell and tissue cultures to aquatic invertebrate cell lines: An updated overview, in: Ballarin, L., Rinkevich, B., Hobmayer, B. (Eds.), Advances in Aquatic Invertebrate Stem Cell Research. MDPI, Basel, pp. 1–64.

Dyachuk, V., 2013. Extracellular matrix is required for muscle differentiation in primary cell cultures of larval *Mytilus trossulus* (Mollusca: Bivalvia). Cytotechnology 65, 725–735. 10.1007/s10616-013-9577-z

Eisenhauer, N., Hines, J., 2021. Invertebrate biodiversity and conservation. Curr. Biol. 31, R1214–R1218. 10.1016/j.cub.2021.06.058

Elagoz, A.M., Van Dijck, M., Lassnig, M., Seuntjens, E., 2024. Embryonic development of a centralised brain in coleoid cephalopods. Neural Dev. 19, 8. 10.1186/s13064-024-00186-2

EUMOFA, European Commission: Directorate-General for Maritime Affairs and Fisheries, E., 2021. Octopus in the EU – Price structure in the supply chain – Focus on Italy, Spain and Greece – Case study. Publications Office. doi10.2771/87203

FAO, 2024. GLOBEFISH - Quarterly cephalopods analysis. Rome.

Fernández-Gago, R., Molist, P., Anadón, R., 2012. Tissues of paralarvae and juvenile cephalopods, in: Gestal, C., Pascual, S., Guerra, A., Fiorito, G., Vieites, J.M. (Eds.), Regeneration and Healing. Springer Open, Gewerbestrasse, pp. 87–112. 10.1007/978-3-030-11330-8

Fiorito, G., Affuso, A., Basil, J., Cole, A., de Girolamo, P., D’angelo, L., Dickel, L., Gestal, C., Grasso, F., Kuba, M., Mark, F., Melillo, D., Osorio, D., Perkins, K., Ponte, G., Shashar, N., Smith, D., Smith, J., Andrews, P.L., 2015. Guidelines for the care and welfare of cephalopods in research –A consensus based on an initiative by CephRes, FELASA and the Boyd Group. Lab. Anim. 49, 1–90. 10.1177/0023677215580006

Flash, T., Zullo, L., 2023. Biomechanics, motor control and dynamic models of the soft limbs of the octopus and other cephalopods. J. Exp. Biol. 226, jeb245295. 10.1242/jeb.245295

García-Fernández, P., Prado-Alvarez, M., Nande, M., Garcia de la serrana, D., Perales-Raya, C., Almansa, E., Varó, I., Gestal, C., 2019. Global impact of diet and temperature over aquaculture of *Octopus vulgaris* paralarvae from a transcriptomic approach. Sci. Rep. 9, 10312. 10.1038/s41598-019-46492-2

Gleadall, I.G., Villanueva, R., Barord, G.J., Doubleday, Z., Aguado-Giménez, F., Akiyama, N., Almansa, E., Ames, C.L., Arkhipkin, A., Avendaño, O., Barrett, C., Bello, G., Bower, J.R., Braga, R., de Luna Sales, J.B., Briceño, F.A., Bustamante, P., Caamal-Monsreal, C., Caballenas-Reboredo, M., Carrasco, S.A., Castellanos-Martínez, S., Cerezo-Valverde, J., Che, L.J., Chung, W., Dan, S., Díaz-Santana-Iturrios, M., Domingues, P., Durante, E.D., Escánez, A., Fernández-Alvárez, F.A., Ferreiro-Velasco, P., Fiorito, G., Furuya, H., Gallardo, P., Ganias, K., Gestal, C., Golikov, A. V., González, A.F., González-Gómez, R., Gordon, J., Guerra, A., Guerrero-Kommritz, J., Hall, K., Haimovici, M., Hamasaki, K., Hernández-Urcera, J., Hirohashi, N., Hirota, K., Hutchinson, N., Imperadore, P., Iwata, Y., Kato, Y., Katugin, O.N., Kimbara, R., Lajbner, Z., Lau, G., Jiménez-Badillo, M.L., Makaida, U., Marquez, L., Mascaro, M., Moltschaniwskyj, N., Morales, A., Moreno, A., Morillo-Velarde, P.S., Nabhitabhata, J., Nande, M., Nishitani, G., Nishshanka, H., Ogura, A., Ortega, A., Ortiz, N., Otero, J., Oyanedel, R., Pang, Y., Pascual, C., Perales-Raya, C., Figueiredo Pereira, J.M., Pita, C., Ponte, G., Power, A.M., Putra, D., Quetglas, A., Repolho, T., Robin, J.P., Rocha, F., Rosa, R., Rosas, C., Rosas-Luis, R., Roumbedajis, K., Roura, Á., Sabirov, R., Sato, N., Sauer, W.H.H., Shaw, P.W., Shigeno, S., De Silva-Dávila, R., Sugimoto, C., Tsukahara, Y., Valls, M., Van der Molen, S., Velázquez-Abunader, I., Vilarnau, D.G., Xavier, J.C., Yoshida, M., Zhang, X., Zheng, J., Zheng, X., Zoral, M.A., 2025. A balanced approach to the potential of octopus aquaculture. Mar. Policy. Accepted.

González, M., Martín-Ruíz, I., Jiménez, S., Pirone, L., Barrio, R., Sutherland, J.D., 2011. Generation of stable *Drosophila* cell lines using multicistronic vectors. Sci. Rep. 1. 10.1038/srep00075

Goswami, M., Yashwanth, B.S., Trudeau, V., Lakra, W.S., 2022. Role and relevance of fish cell lines in advanced in vitro research. Mol. Biol. Rep. 49, 2393–2411. 10.1007/s11033-021-06997-4

Haslam, G., Wyatt, D., Kitos, P.A., 2000. Estimating the number of viable animal cells in multi-well cultures based on their lactate dehydrogenase activities. Cytotechnology 32, 63–75. 10.1023/A:1008121125755

Hendijani, F., 2017. Explant culture: An advantageous method for isolation of mesenchymal stem cells from human tissues. Cell Prolif. 50, e12334. 10.1111/cpr.12334

Jacquet, J., Franks, B., Godfrey-Smith, P., Sánchez-Suárez, W., 2019. The case against octopus farming. Issues Sci. Technol. 35, 37–44. 10.2307/26948988

Kim, Y., Tanner, H.M., Rosenthal, J.J.C., Brangwynne, C.P., 2025. Squid primary cell culture as a model system and experimental tool. bioRxiv. 10.1101/2025.01.12.632648

Kitaeva, K.V., Rutland, C.S., Rizvanov, A.A., Solovyeva, V. V., 2020. Cell culture based in vitro test systems for anticancer drug screening. Front. Bioeng. Biotechnol. 8, 322. 10.3389/fbioe.2020.00322

Leelavatcharamas, V., Emery, A.N., Al-Rubeai, M., 2020. Monitoring the proliferative capacity of cultured animal cells by cell cycle analysis, in: Al-Rubeai, M., Emery, A.N. (Eds.), Flow Cytometry Applications in Cell Culture. CRC Press, Boca Ratón, pp. 1–15. 10.1201/9781003067467

Lordon, B., Campion, T., Gibot, L., Gallot, G., 2024. Impact of trypsin on cell cytoplasm during detachment of cells studied by terahertz sensing. Biophys. J. 123, 2476–2483. 10.1016/j.bpj.2024.06.011

Lyons, A.B., Parish, C.R., 1994. Determination of lymphocyte division by flow cytometry. J. Immunol. Methods 171, 131–137. 10.1016/0022-1759(94)90236-4

Maiorova, M.A., Odintsova, N.A., 2016. Proliferative potential of larval cells of the mussel *Mytilus trossulus* and their capacity to differentiate into myogenic cells in culture. Russ. J. Mar. Biol. 42, 281–285. 10.1134/S1063074016030068

Malham, S.K., Runham, N.W., Secombes, C.J., 1998. Lysozyme and antiprotease activity in the lesser octopus *Eledone cirrhosa* (Lam.) (Cephalopoda). Dev. Comp. Immunol. 22, 27–37. 10.1016/S0145-305X(97)00045-1

Márquez, L., Martín, M.V., Larson, M., Almansa, E., 2023. Application of the thermal time model of embryonic development duration to the culture of octopod cephalopods. Aquac. Reports 30, 101563. 10.1016/j.aqrep.2023.101563

Maselli, V., Xu, F., Syed, N.I., Polese, G., Di Cosmo, A., 2018. A novel approach to primary cell culture for *Octopus vulgaris* neurons. Front. Physiol. 9, 220. 10.3389/fphys.2018.00220

Morales, A.E., Cardenete, G., Hidalgo, M.C., Garrido, D., Martín, M.V., Almansa, E., 2017. Time course of metabolic capacities in paralarvae of the common octopus, Octopus vulgaris, in the first stages of life. searching biomarkers of nutritional imbalance. Front. Physiol. 8, 427. 10.3389/fphys.2017.00427

Mótyán, J., Tóth, F., Tőzsér, J., 2013. Research applications of proteolytic enzymes in molecular biology. Biomolecules 3, 923–942. 10.3390/biom3040923

Mukherjee, T.K., Malik, P., Mukherjee, S., 2023. Common reagents and medium for mammalian cell culture, in: Mukherjee, T.K., Malik, P., Mukherjee, S. (Eds.), Practical Approach to Mammalian Cell and Organ Culture. Springer Nature, Singapore, pp. 1–158. 10.1007/978-981-19-1731-8

Nesher, N., Maiole, F., Shomrat, T., Hochner, B., Zullo, L., 2019. From synaptic input to muscle contraction: Arm muscle cells of *Octopus vulgaris* show unique neuromuscular junction and excitation-contraction coupling properties. Proc. R. Soc. B Biol. Sci. 286. 10.1098/rspb.2019.1278

Odintsova, N.A., Dyachuk, V.A., Nezlin, L.P., 2010. Muscle and neuronal differentiation in primary cell culture of larval *Mytilus trossulus* (Mollusca: Bivalvia). Cell Tissue Res. 339, 625–637. 10.1007/s00441-009-0918-3

Odintsova, N.A., Plotnikov, S.V., Karpenko, A.A., 2000. Isolation and partial characterization of myogenic cells from mussel larvae in vitro. Tissue Cell 32, 417–424. 10.1054/tice.2000.0130

Pita, C., Roumbedakis, K., Fonseca, T., Matos, F.L., Pereira, J., Villasante, S., Pita, P., Bellido, J.M., Gonzalez, A.F., García-Tasende, M., Lefkaditou, E., Adamidou, A., Cuccu, D., Belcari, P., Moreno, A., Pierce, G.J., 2021. Fisheries for common octopus in Europe: socioeconomic importance and management. Fish. Res. 235, 105820. 10.1016/j.fishres.2020.105820

Ponte, G., Roumbedakis, K., Galligioni, V., Dickel, L., Bellanger, C., Pereira, J., Vidal, E.A.G., Grigoriou, P., Alleva, E., Santucci, D., Gili, C., Botta, G., Imperadore, P., Tarallo, A., Juergens, L., Northrup, E., Anderson, D., Aricò, A., De Luca, M., Pieroni, E.M., Fiorito, G., 2023. General and species-specific recommendations for minimal requirements for the use of cephalopods in scientific research. Lab. Anim. 57, 26–39. 10.1177/00236772221111261

Prado-Álvarez, M., Dios, S., García-Fernández, P., Tur, R., Hachero-Cruzado, I., Domingues, P., Almansa, E., Varó, I., Gestal, C., 2022. De novo transcriptome reconstruction in aquacultured early life stages of the cephalopod Octopus vulgaris. Sci. Data 9, 609. 10.1038/s41597-022-01735-2

Reis, D.B., Shcherbakova, A., Riera, R., Martín, V.M., Domingues, P., Andrade, J.P., Jiménez-Prada, P., Rodríguez, C., Sykes, A. V, Almansa, E., 2021. Effects of feeding with different live preys on the lipid composition, growth and survival of *Octopus vulgaris* paralarvae. Aquac. Res. 52, 105–116. 10.1111/are.14873

Rinkevich, B., 1999. Cell cultures from marine invertebrates: obstacles, new approaches and recent improvements. Prog. Ind. Microbiol. 35, 133–153. 10.1016/s0168-1656(99)00067-x

Romano, G., Almeida, M., Coelho, A.V., Cutignano, A., Gonçalves, L.G., Hansen, E., Khnykin, D., Mass, T., Ramšak, A., Rocha, M., da Silva, T.H., Sugni, M., Ballarin, L., Geneviere, A.-M., 2022. Bioactive natural products and biomaterials from marine inver3 tebrates: from basic research to innovative applications. Manuscript submitted for publication. Mar. Drugs 20, 1–45. 10.3390/md20040219

Rosa, R., Santos, C.P., Borges, F., Amodio, P., Amor, M., Bower, J.R., Caldwell, R.L., Di Cosmo, A., Court, M., Fiorito, G., Gestal, C., González, Á.F., Guerra, Á., Hanlon, R.T., Hofmeister, J.K.K., Ibáñez, C.M., Ikeda, Y., Imperadore, P., Kommritz, J.G., Kuba, M., Hall, K.C., Lajbner, Z., Leite, T.S., Lopes, V.M., Markaida, U., Moltschaniwskyj, N.A., Nabhitabhata, J., Ortiz, N., Otjacques, E., Pizzulli, F., Ponte, G., Polese, G., Raffini, F., Rosas, C., Roura, Á., Sampaio, E., Segawa, S., Simakov, O., Sobrino, I., Storero, L.P., Voight, J.R., Williams, B.L., Zheng, X., Pierce, G.J., Villanueva, R., Gleadall, I.G., 2024. Chapter 23 - Past, present, and future trends in octopus research, in: Rosa, R., Gleadall, I.G., Pierce, G.J., Villanueva, R.B.T.-O.B. and E. (Eds.),. Academic Press, pp. 421–454. 10.1016/B978-0-12-820639-3.00010-8

Sauer, W.H.H., Gleadall, I.G., Downey-Breedt, N., Doubleday, Z., Gillespie, G., Haimovici, M., Ibáñez, C.M., Katugin, O.N., Leporati, S., Lipinski, M.R., Markaida, U., Ramos, J.E., Rosa, R., Villanueva, R., Arguelles, J., Briceño, F.A., Carrasco, S.A., Che, L.J., Chen, C.S., Cisneros, R., Conners, E., Crespi-Abril, A.C., Kulik, V.V., Drobyazin, E.N., Emery, T., Fernández-Álvarez, F.A., Furuya, H., González, L.W., Gough, C., Krishnan, P., Kumar, B., Leite, T., Lu, C.C., Mohamed, K.S., Nabhitabhata, J., Noro, K., Petchkamnerd, J., Putra, D., Rocliffe, S., Sajikumar, K.K., Sakaguchi, H., Samuel, D., Sasikumar, G., Wada, T., Zheng, X., Tian, Y., Pang, Y., Yamrungrueng, A., Pecl, G., 2021. World Octopus Fisheries. Rev. Fish. Sci. Aquac. 29, 279–429. 10.1080/23308249.2019.1680603

Seibel, B.A., Childress, J.J., 2000. Metabolism of benthic octopods (Cephalopoda) as a function of habitat depth and oxygen concentration 47, 1247–1260. 10.1016/S0967-0637(99)00103-X

Shen, Y., Vignali, P., Wang, R., 2017. Rapid profiling cell cycle by flow cytometry using concurrent staining of DNA and mitotic markers. Bio-Protocol 7, 2–5. 10.21769/bioprotoc.2517

Slanzi, A., Iannoto, G., Rossi, B., Zenaro, E., Constantin, G., 2020. *In vitro* models of neurodegenerative diseases. Front. Cell Dev. Biol. 8, 328. 10.3389/fcell.2020.00328

Tur, R., Domingues, P., Almansa, E., Lago, M., García-Fernández, P., Pérez, E., 2020. Procedimiento para el cultivo de paralarvas del pulpo común Octopus vulgaris. ES2714930.

Van Der Merwe, M., Auzoux-Bordenave, S., Niesler, C., Roodt-Wilding, R., 2010. Investigating the establishment of primary cell culture from different abalone (*Haliotis midae*) tissues. Cytotechnology 62, 265–277. 10.1007/s10616-010-9293-x

Yoshino, T.P., Bickham, U., Bayne, C.J., 2013. Molluscan cells in culture: primary cell cultures and cell lines. Can. J. Zool. 91, 391–404. 10.1139/cjz-2012-0258

Zhang, S., Kuhn, J.R., 2013. Cell isolation and culture. WormBook 1–39. 10.1895/wormbook.1.157.1

